## Supplemental Material 1 for "Origin of imported SARS-CoV-2 strains in The Gambia identified from Whole Genome Sequences"

**Method:**

**Sample acquisition**

NPS samples from COVID-19 positive patients were requested from the diagnostic team for genomic sequencing. A total of three submissions were made, first submission involved two samples from the first COVID-19 positive patient and the second was nine samples and the third involved two samples as detailed in table 1.

**RNA extraction**

Total RNA was purified from thirteen samples (details in table 1) using the QiaAmp RNA mini kit (Qiagen – 52906) following viral inactivation at our MRCG containment level 3 facility. The purified total RNA samples were quantified using the high sensitivity dsDNA Qubit reagent kit on Qubit fluorometer 3.0 and RNA integrity was checked on the Agilent Tapestation 4200.

Table 1: Sample information for COVID-19 sequenced cases from The Gambia

| **Case ID** | **Age (yrs)** | **Sex** | **Travelled from** | **Date Reported** | **Current Status** | **Number of samples submitted** | **Time points** | **Library prep type** | | | **Sequencing** | |
| --- | --- | --- | --- | --- | --- | --- | --- | --- | --- | --- | --- | --- |
|  |  |  |  |  |  |  |  | **Depletion** | **ARTIC amplicon (NEB)** | **ARTIC amplicon (ONT -LSK109)** | **Illumina (MiSeq)** | **Nanopore (GridION)** |
| A | 28 | F | London | 16/03/20 | Recovered | 4 | Days 0,4,7,10 | 2 | 4 | 4 | 4 | 4 |
| B | 70 | M | Bangladesh | 19/03/20 | Dead | 1 | Day 0 | 0 | 1 | 1 | 1 | 1 |
| C | 71 | M | France | 20/03/20 | Recovered | 1 | Day 0, | 0 | 1 | 1 | 1 | 1 |
| D | 53 | M | France | 26/03/20 | Active | 2 | Day 0,11 | 0 | 1 | 2 | 1 | 2 |
| E | 21 | F | Netherland | 23/03/20 | Active | 2 | Day 0,14 | 0 | 1 | 2 | 1 | 2 |
| F | 30 | M | Italy | 13/03/20 | Recovered | 1 | Day 0 | 0 | 1 | 1 | 1 | 1 |
| Total | | | | | | 11 |  | 2 | 9 | 11 | 9 | 11 |
| Cases A-D = Confirmed RT-PCR COVID-19 cases | | | | |  |  |  |  |  |  |  |  |
| Case E = Indeterminate by RT-PCR | | | | |  |  |  |  |  |  |  |  |
| Case F = RT-PCR COVID-19 negative | | | | |  |  |  |  |  |  |  |  |
| Cases A and D travelled to The Gambia in the same flight | | | | |  |  |  |  |  |  |  |  |
| Cases C and D both travelled from France | | | | |  |  |  |  |  |  |  |  |

**Ribosomal RNA depletion for Batch 1 submission**

The two samples were depleted using the RiboMinus transcription isolation kit from ThermoFisher and purified using RNA purification beads from Beckman Coulter.

**cDNA conversion**

The purified RiboMinus samples were converted to cDNA as per the NEBNext ultra II RNA library prep kit (NEB, E7770L). Total RNA from batch 2 and 3 submissions were converted to cDNA as per the ARTIC amplicon sequencing protocol for nCoV-2019^1^ by adding 1μl of 50μM random hexamers (NEB) and 1μl of 10mM dNTPs mix to 11μl of the purified RNA. These were incubated in a thermocycler at 65°C for 5 minutes and cooled on ice for 1 minute. The annealed template RNA was combined with 4μl SSIV buffer, 1μl 100mM DTT, 1μl RNaseOUT RNase Inhibitor and 1μl SSIV Reverse Transcriptase and incubated at 42°C for 50 minutes and 70°C for 10 minutes to synthesise the second strand.

**Multiplex PCR**

Version 2 of the primer schemes for nCoV-2019 were ordered from Metabion international and reconstituted as per the ARTIC protocol^1^. Two primer pools at 10μM containing 98 primers each were used for the PCR amplification. For each sample, two mastermixes were prepared; one containing primer pool1 and the other primer pool2. Template cDNA (2.5μl) was added to a mastermix containing the following components; 5μl 5X Q5 reaction buffer, 0.5μl 10mM dNTPs, 0.25μl Q5 Hot start DNA polymerase, 3.6μl of primer pool1 and 2 and 13.15μl of nuclease free water. The tubes were mixed and centrifuged before being subjected to the following cycling condition: heat activation at 98°C for 30 seconds, denaturation at 98°C for 15 seconds and annealing at 65°C for 5 minutes. The denaturation and annealing steps were repeated for a total of 35 cycles. The PCR products of pool1 and 2 were combined and purified using Ampure beads (Beckman Coulter) as per the ARTIC protocol. The purified products were visualised using the Agilent Tapestation 4200 and quantified using the Qubit fluorometer (Invitrogen).

**Illumina and Nanopore library Preparation and sequencing**

**Illumina**

The purified cDNA from the depletion and PCR products from the ARTIC protocol were normalised to 100ng with EB buffer (10mM Tris-HCl) to a final volume of 25µl for Illumina library preparation using the NEBNext ultra II DNA library prep kit for Illumina (New England Biolabs, UK; E7645). The Amplicons were end repaired by adding 3.5µl of the end repair buffer and 1.5µl of the end repair enzyme and incubated in a thermocycler at 20^°^C for 30min and 65^°^C for 30min with the heated lid set to ≥75^°^C. The end repaired products were adaptor ligated by adding 15µl of the NEBNext ligation mix, 0.5µl of the ligation enhancer and 1.25µl of 10-fold diluted adaptor and incubated in a thermocycler for 15mins at 20^°^C with the heated lid off. The ligated fragments were digested by adding 1.5µl of user enzyme and incubated in a thermocycler for 15mins at 37^°^C with the lid set to 47^°^C. The digested ligated products were purified using Ampure XP beads (Beckman Coulter A63881) according to the NEBNext® Ultra™ II DNA Library Prep Kit for Illumina® protocol (E7645) (New England Biolabs, UK). The purified products were PCR enriched by adding 12.5µl of the Q5 mastermix, 2.5µl of the i5 and i7 indexes to the 7.5µl elute from the purification. The samples were placed in a thermocycler and amplified for 7 cycles at 98^°^C for 30sec, 98^°^C for 10sec, 65^°^C for 75sec, 65^°^C for 5mins and held at 4^°^C. Following the PCR, the samples were purified and quantified using the high sensitivity dsDNA Qubit kit and sized using D1000 ScreenTape on the Agilent Tapestation 4200. Molarity was calculated using size and concentration and all libraries were normalised to 10nM before pooling. The final pool was quantified on a Qubit 3.0 instrument and run on D1000 ScreenTape (Agilent Catalogue No. 5067-5579) using the Agilent Tapestation 4200 to calculate the final library pool molarity.

The pool was run at a final concentration of 10pM on an Illumina Miseq instrument using Miseq V3 reagent kit 600 cycles following the Illumina recommended denaturation and loading recommendations which included a 5% PhiX spike in (PhiX Control v3 Illumina Catalogue FC-110-3001). Data was uploaded to BaseSpace (www.basespace.illumina.com) where the raw data was converted to FASTQ files for each sample.

**Nanopore**

Library preparation on the purified PCR product from the ARTIC protocol was done using the Nanopore ligation sequencing kit (SQK-LSK109). The adaptor-ligated libraries were quantified on Qubit Fluorometer 3.0 and sized on Agilent Tapestation 4200 before loading 20nM onto a primed GridION flow cell for sequencing. The sequences were concurrently run on RAMPART, which provides real-time overview of genome coverage as well as reference matching for each sample.

Table 1a. Reference Strains

| Name | Accession Number | Country |
| --- | --- | --- |
| SARS-CoV-2 | MT292569.1 | Spain |
| SARS-CoV-2 | MT007544.1 | Australia – Victoria |
| SARS-CoV-2 | MT050493.1 | India – Kerala |
| SARS-CoV-2 | MT066156.1 | Italy |
| SARS-CoV-2 | LC528223.1 | Japan |
| SARS-CoV-2 | MN985325.1 | USA – Washington |
| SARS-CoV-2 | MT039890.1 | South Korea |
| SARS-CoV-2 | MN908947.3 | China - Wuhan |
